# The paradox of extremely fast evolution driven by genetic drift in multi-copy gene systems

**DOI:** 10.1101/2023.06.14.545040

**Authors:** Xiaopei Wang, Yongsen Ruan, Lingjie Zhang, Xiangnyu Chen, Bingjie Chen, Miles Tracy, Liying Huang, Zhongqi Liufu, Chung-I Wu, Haijun Wen

**Author notes:** Corresponding authors (H. Wen); (C.I. Wu).

## Abstract

Multi-copy gene systems that evolve within, as well as between, individuals are common. They include viruses, mitochondrial DNAs, multi-gene families etc. The paradox is that neutral evolution in two stages should be slower than single-copy systems but the opposite is often true. We now apply the Haldane model, recently generalized as the GH model (1), to quantify genetic drift in mammalian ribosomal RNA genes (rDNAs). On average, the copy number (*C*) is 150 - 300 per haploid. A neutral mutation in rDNA should take 4*NC*^*\**^ generations to become fixed (*N*, the population size; *C* ^*\**^, the effective copy number within individuals). While *C* > *C*^*\**^ >> 1 is expected, the observed fixation time in mouse is < 4*N*, leading to the paradox of *C*^*\**^ < 1. Genetic drift of rRNA genes thus appears 10-100 times stronger than in single-copy genes. The large increases are driven by a host of molecular mechanisms such as gene conversion and unequal crossover. Although each mechanism of drift is very difficult to quantify, the GH model permits the estimation of their total effects that constitute the aggregate “evolutionary noises”. In humans, the fixation rate of rRNA genes is higher than the theoretical maximum of drift, hence, justifying the inference of adaptive evolution. In conclusion, the stochastic evolution in multi-copy gene systems, including viruses and others, can be effectively tracked by GH model.

## Introduction

In molecular evolution, random forces are rampant, but the standard Wright-Fisher (WF) model cannot account for many of them, thus leading to the conflation of random noises with adaptation (1)(the companion submission, Ruan et al. (2025)). In this study, we focus on multi-copy gene systems whereby the evolution takes place in two stages: both within (stage I) and between individuals (stage II). Multi-copy gene systems include viruses, transposons, mitochondria and multi-gene families. Given the extra stage of within-host fixation, the neutral evolutionary rate of multi-copy systems should be much slower than in single-copy systems. However, the rapid evolution of multi-copy systems has been extensively documented (2–5).

The speed of neutral evolution is the basis for analyzing molecular evolution. Neutral evolution is driven by random transmission, gene conversion, stochastic replication etc., which collectively constitute genetic drift. All other evolutionary forces, such as selection, mutation and migration, can be analyzed only after genetic drift fully is accounted for. The standard Wright-Fisher (WF) model of genetic drift tracks neutral evolution by the random sampling of genes in a population of size *N*. For deviations from the ideal population, many modifications of the WF model (6–10) have been proposed that substitute various *N*_*e*_’s (effective population size) for the actual *N*.

The companion paper (1) presents three paradoxes of genetic drift inherent in the WF models. A most curious one is that genetic drift may often become stronger as *N* increases (11–13). Here, we focus on the fourth paradox that arises in multi-copy gene systems that include viral epidemics (14), transposons (15), mitochondrial DNAs (16), satellite DNAs (17, 18), and ribosomal RNA genes (19, 20). In COVID-19, the inability of the WF models to track both within- and between-host evolution simultaneously is a main reason for much confusion about the origin, spread and driving forces of SARS-CoV-2 (2, 21–23).

Ruan et al. (2025) re-introduces the Haldane model of the branching process to account for random forces in molecular evolution. It is generalized to track changes in population size. Here, we apply the GH (Generalized Haldane) model to multi-copy gene systems, using ribosomal RNA genes (rDNAs) as an example. In such systems, the molecular mechanisms driving neutral evolution within individuals (Stage I) are diverse. The literature has shown the difficulty in modeling each molecular mechanism specifically (16, 24–30). These mechanisms are also distinct from the mechanisms in Stage II, making the estimation of the total genetic drift most challenging.

The GH model does not analyze each molecular mechanism of drift individually. Instead, it filters out their total effect in the form of variance in reproductive output (or *V(K)*). By doing so, the GH model can effectively track genetic drift in multi-copy gene systems.

## Results

PART I presents background analyses of the rRNA genes. PART II consolidates aspects of the GH model for analyzing the rDNA polymorphisms and divergence data. In PART III, we apply the theory to rRNA evolution in mice and apes.

### PART I - The pseudo-population of ribosomal RNA genes within each individual

The ribosomal RNA genes (or simply rDNAs) are multi-copy gene clusters (31) that are arrayed as tandem repeats on multiple chromosomes as shown in Fig. 1A (32, 33). In humans, the copy number can vary from 60 to 1600 per individual (mean, 315; SD, 104; median, 301) (34). For each haploid genome, *C* ~ 150 on average in humans and *C* ~ 110 in mice (34). In humans, the five rRNA clusters are located on the short arm of the five acrocentric chromosomes (35). Such an arrangement permits crossovers between chromosomes without perturbing the rest of the genomes. In *Mus*, the rDNAs are all located in the pericentromeric or sub-telomeric region, on the long arms of telocentric chromosomes (33, 36). Thus, unequal crossovers between non-homologous chromosomes may involve centromeres while other genic regions are also minimally perturbed.

**Figure 1.**
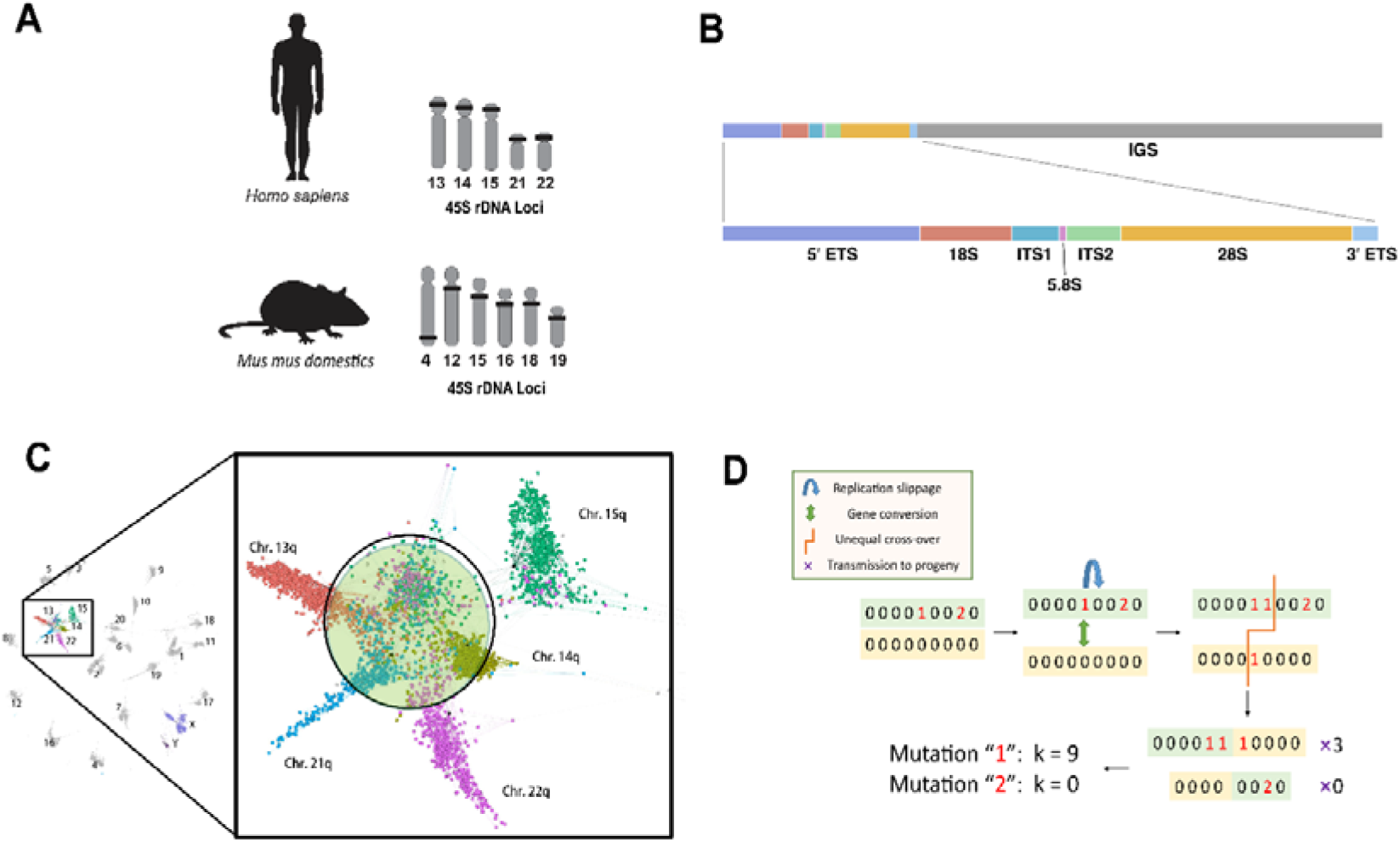
The “chromosome community of rDNAs on five acrocentric chromosomes. (A) The genomic locations of rDNA tandem repeats in human (70) and mouse (33). rDNAs are located on the short arms (human), or the proximal end of the long arms (mouse), of the chromosome. Either way, inter-chromosomal exchanges are permissible. (B) The organization of rDNA repeat unit. IGS (intergenic spacer) is not transcribed. Among the transcribed regions, 18S, 5.8S and 28S segments are in the mature rRNA while ETS (external transcribed spacer) and ITS (internal transcribed spacer) are excluded. (C) The pseudo-population of rRNA genes is shown by the “chromosomes community” map (43), which indicates the divergence distance among chromosome segments. The large circle encompasses rDNAs from all 5 chromosomes. It shows the concerted evolution among rRNA genes from all chromosomes, which thus resemble members of a (pseudo-)population. The slightly smaller thin circle, from the analysis of this study, shows that the rDNA gene pool from each individual captures approximately 95% of the total diversity of human population. (D) A simple illustration that shows the transmissions of two new mutations (#1 and #2 in red letter). Mutation 1 experiences replication slippage, gene conversion and unequal crossover and grows to 9 copies (*K* = 9) after transmission. Mutation 2 emerges and disappears (*K* = 0). This shows how *V(K)* may be augmented by the homogenization process.

Each copy of rRNA gene has a functional and non-functional part as shown in Fig. 1B. The “functional” regions of rDNA, 18S, 5.8S, and 28S rDNA, are believed to be under strong negative selection, resulting in a slow evolution rate in animals (37). In contrast, the transcribed spacer (ETS and ITS) and the intergenic spacer (IGS) are much less constrained by negative selection and often labeled “non-functional” (4). In this study of genetic drift, we focus on the non-functional parts.

While a human haploid with 200 rRNA genes may appear to have 200 loci, the concept of “gene loci” cannot be applied to the rRNA gene clusters. This is because DNA sequences can spread from one copy to others on the same chromosome via replication slippage. They can also spread among copies on different chromosomes via gene conversion and unequal crossovers (25, 26, 35, 38). These mechanisms will be referred to collectively as the homogenization process. Copies of the cluster on the same chromosome are known to be nearly identical in sequences (20, 39). There is also extensive evidence for genetic exchanges between chromosomes (40–42).

In short, rRNA gene copies in an individual can be treated as a pseudo-population, corresponding to the recently proposed “chromosome community” (43). A communal pool of rRNA genes is distributed among the short arms of the five acrocentric chromosomes (Fig. 1C). These copies can move within or between chromosomes by the mechanisms depicted in Fig. 1D. The entire human population is thus a collection of sub-populations, each of which is a pseudo-population of rRNA genes within an individual.

### PART II - Theory

#### 1. The Haldane model of genetics drift applied to multi-copy gene systems

The Haldane model of genetic drift (44) is based on the branching process. It is a powerful alternative to the Wright-Fisher model (29, 45, 46). In the Haldane model, each copy of the gene leaves *K* copies in a time interval with the mean and variance of *E*(*K*) and *V*(*K*). If *V*(*K*) = 0, there is no gene frequency change and no genetic drift. In the standard WF model, *V*(*K*) = *E*(*K*), which has been relaxed in later modifications (6–10, 45).

In the companion study, Ruan et al. (2025) propose a Generalized Haldane (GH) model that incorporateds changes in the population size (*N*). The GH model thus tracks relative gene frequencies as well as absolute copy numbers. Here, we compare the relative strength of genetic drift in rRNA genes vs. single-copy genes using the GH model. We shall use * to designate symbols for rRNA genes. In this study, both *E*(*K*) and *E*^*\**^(*K*) are set to 1 such that the long-term copy number in the population remains constant.

In the GH model, genetic drift is determined by *N/V(K)* for single copy genes. (The notion of *N*_*e*_ = *N/V(K)* of the WF model is not adopted in the GH model.) For simplicity, we assume *V(K)* = 1 for single-copy genes.

For rRNA genes, *V*^*\**^(*K*) ≥ 1 may generally be true because *K* for rDNA mutations are affected by a host of random homogenization factors including replication slippage, unequal cross-over, gene conversion and other related mechanisms not operating on single-copy genes (Fig. 1D). Hence, genetic drift in rDNA is determined by

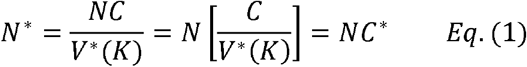

where *C* is the average number of rRNA genes in an individual and *V*^*\**^*(K)* reflects the homogenization process on rRNA genes. Thus,

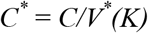

represents the effective copy number of rRNA genes in the population. *C*^*\**^ determines the strength of genetic drift in rRNA genes relative to single-copy genes. Since *C* is in the hundreds and *V*^*\**^*(K)* is expected to be > 1, the relationship of 1 << *C*^*\**^≤*C* is hypothesized. Fig. 1D is a simple illustration that the homogenizing process may enhance *V*^*\**^(*K*) substantially over the WF model.

#### 2. rDNA vs. single-copy gene polymorphism within species

Genetic drift is manifested in the level of heterozygosity (*H*), the probability that two randomly chosen genes from the population are of different allelic classes. *H* is proportional to *N* (or *N*^*\**^) when *H* is small. Therefore, *H* should be larger for rRNA genes than for single-copy genes (by > 5 fold in both mouse and human; see PART III).

For rRNA genes, in addition to the *H* value of the species, *H* can also be the heterozygosity within each individual, to be referred as *H*_*I*_. The population level *H* would then be *H*_*S*_ (see Supplementary Note 1 for details). An issue of interest is the differentiation of rRNA genes among individuals, defined as

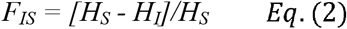

where *H*_*I*_ would be averaged over all individuals. With a low *F*_*IS*_ value, say, < 0.1, the diversity of rRNA genes in a single individual would capture > 90% of the diversity of the entire species.

#### 3. rDNA divergence between species

Whereas the level of genetic diversity is a function of the population size, the rate of divergence between species, in its basic form, does not depend on *N*. The rate of neutral molecular evolution (λ) is generally shown by Eq. (3) for single-copy genes (47–49):

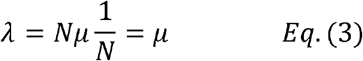

The factor of 1/*N* in Eq. (3) indicates the fixation probability of a new mutation. If we substitute *N*^*\**^ for *N*, Eq. (3) is applicable to the long-term evolutionary rate of rRNA genes as well.

**Figure 2.**
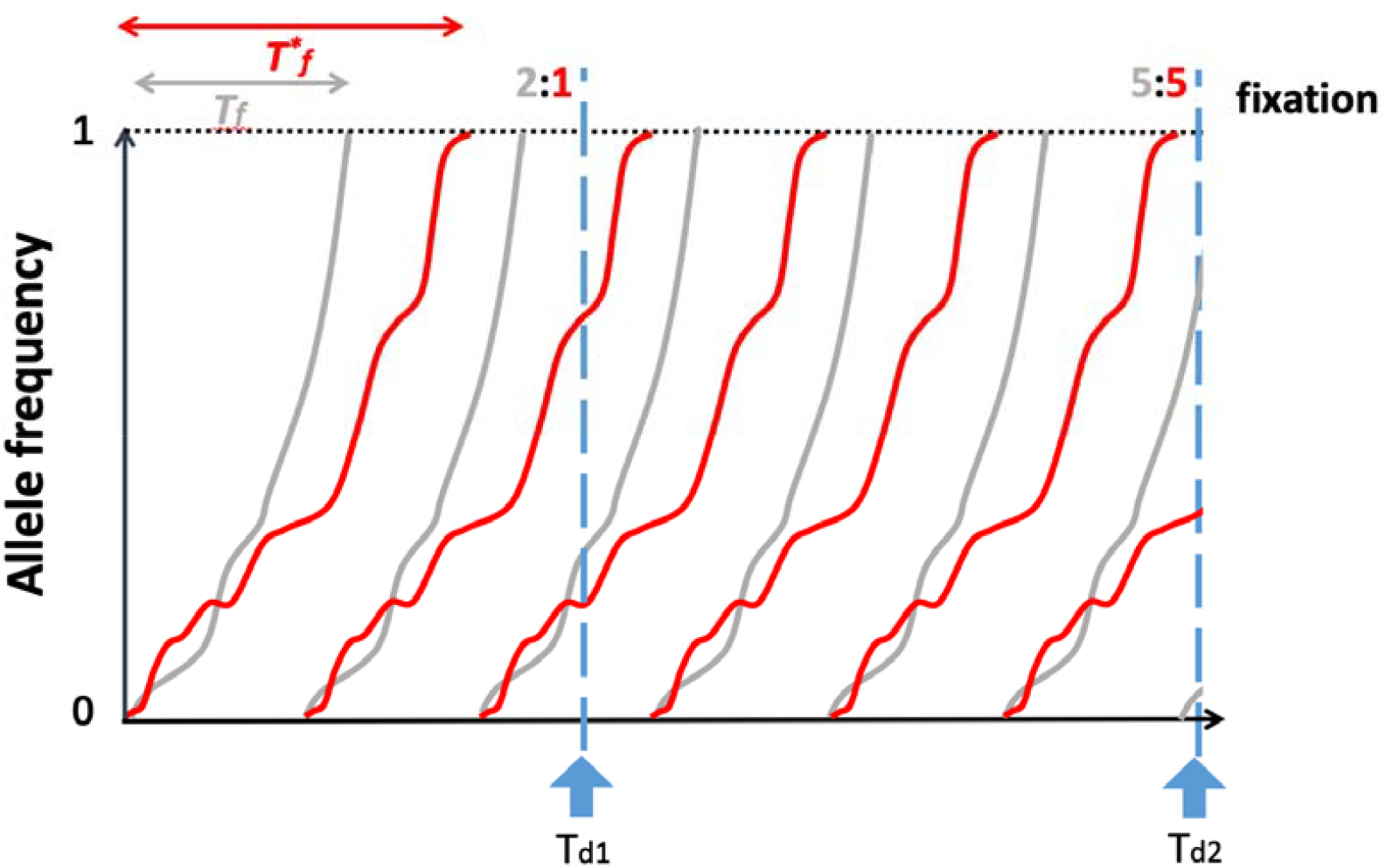
Fixation of mutations between closely related species (at the shorter divergence time of *T*_*d1*_). We assume two classes of mutations with different fixation times - *T*_*f*_ (shorter, grey line) and *T*^*\**^_*f*_ (longer, red line). Specifically, the red lines designate rRNA mutations as the polymorphism data indicate *N*^*\**^ > *N* (see main text). The figure shows that rRNA mutations with a longer *T*^*\**^_*f*_ would have a lower fixation rate at *T*_*d1*_ (2:1) than single-copy genes. In the longer term (*T*_*d2*_), the ratio (5:5) would follow Eq. (3) whereby the fixation time ceases to have an effect.

Eq. (3), however, is for long-term evolution. For shorter-term evolution (at *T*_*d1*_ in Fig. 2), it is necessary to factor in the fixation time (Fig. 2), *T*_*f*_, which is the time between the emergence of a mutation and its fixation. If we study two species with a divergent time (*T*_*d1*_) equal to or smaller than *T*_*f*_, then few mutations of recent emergence would have been fixed as species diverge, thereby limiting the observed divergence. For example, *T*_*d*_ is about 6 million years (Myrs) between human and chimpanzee while *T*_*f*_ is roughly 0.8 Myrs in humans. Mutations would not get fixed during the recent 0.8 Myrs, indicated in Fig. 2. Consequently, the realized substitution rate for single-copy genes would be fraction lower *T*_*f*_ */T*_*d*_ (~ 0.8/6 = 0.13) than Eq. (3) would indicate. This reduction would approach zero as *T*_*d*_ increases. Assuming *C*^*\**^ > 1, Fig. 2 depicts the fixation rate of single copy genes (grey lines) to be higher than rRNA genes (red lines) at *T*_*d1*_. However, as will be shown in PART III.2, this prediction is often not true, implying that *C*^*\**^ < 1. Hence, the paradox is that the effective copy number of rRNA genes is less than one.

### PART III - Data Analyses

#### 1. rDNA polymorphism within species

##### 1) Polymorphism in mice

In this study, rRNA polymorphism is based on the non-functional portion of the rRNA gene which should be close to the neutral diversity. In *M. m. domestic, H*_*I*_ of 10 individuals ranges from 0.00556 to 0.00673 while *H*_*S*_ is 0.00725 (see Table S1). Thus, *F*_*IS*_ = [*H*_*S*_ - *H*_*I*_]/*H*_*S*_ for mice is 0.142, which means that 86% of within-species variation can be found within each individual. Obviouslym there is no *F*_*IS*_ for single copy genes and *H*_*S*_ for rRNA genes is 0.00725, or 7.25 per kb, is 5.2 times larger than that of single copy genes at 1.40 per kb genome-wide (50). In other words, *C*^*\**^ = *N*^*\**^*/N* ~ 5.2. The *H*_*S*_ results confirm the prediction that rRNA genes should be more polymorphic than single-copy genes. Given that each haploid genome has ~ 110 copies of rRNA genes, the 5.2 ratio hints a degree of homogenization.

##### 2) Polymorphism in human

*F*_*IS*_ for rDNA among 8 human individuals is 0.059 (Table S2), even smaller than 0.142 in *M. m. domesticus* mice, indicating minimal genetic differences among human individuals. Consistent with a low *F*_*IS*_, Fig. S1 shows strong correlation of the polymorphic site frequencies of rDNA transcribed and IGS regions among each pair of individuals from three continents (2 Asians, 2 Europeans and 2 Africans). The results suggest that the genetic drift in the population is augmented substantially by the homogenization process within individuals. In humans as in mice, intra-species polymorphism is mainly preserved within individuals.

The heterozygosity (*H*_*S*_) has been reported to be around 0.8 – 0.9 per Kb (51) in contrast with the *H*_*S*_ for rRNA genes at 7.24 per Kb (Table S2). Hence, the effective copy number of rDNAs is roughly, or *C*^*\**^ ~ 8 (7.2 /0.9). This reduction from the actual copy number of 150 to *C*^*\**^ ~ 8 again suggests strong homogenization within individuals.

##### 3) Further evidence of homogenization within species

Fig. 3B and 3C show the distribution of *F*_*IS*_ values among variant sites on rRNA genes. In 3B, *F*_*IS*_ values are generally below 0.2 among three wild *M. m. domesticus* mice. In 3C, the distribution shifts to higher values among 10 mouse strains, seven of which are highly inbred. The differences indicate that inbred strains have experienced strong drift within each line but such strain differentiation disappears among outcross strains. A new peak in Fig. 3C above 0.2 suggests that the speed of homogenization within each inbred line (and hence differentiation between lines) is substantial.

**Figure 3.**
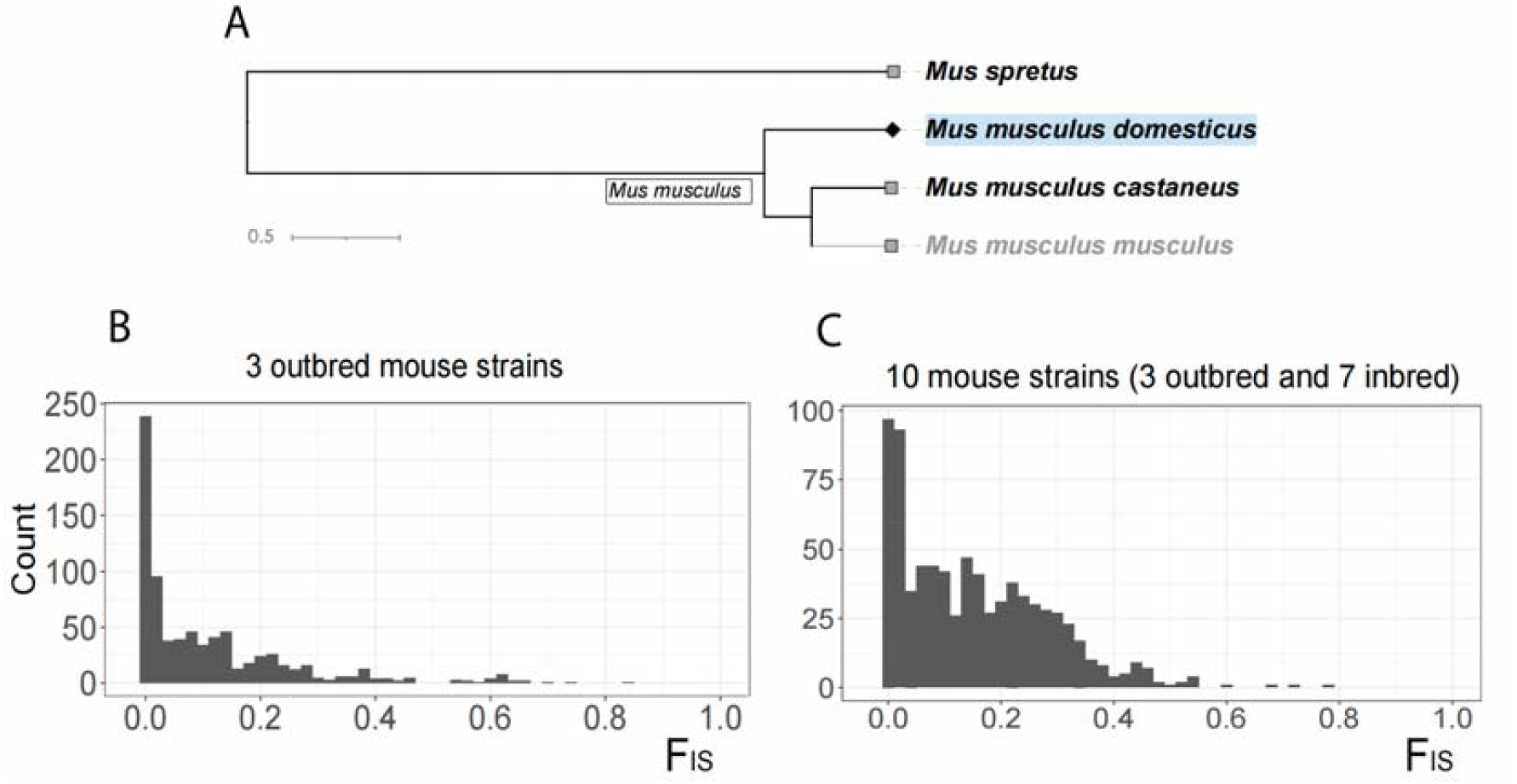
Distribution of rRNA variation within- vs. between individual mice. (A) Phylogeny of *Mus musculus* and *Mus spretus* mice used in this study. The divergence times are obtained from http://timetree.org/. The line segment labeled 0.5 represents 0.5 Myrs. (B, C) *F*_*IS*_ distribution within *M. m. domesticus*. The distributions of *F*_*IS*_ for polymorphic sites in 3 outbred mouse strains and all 10 mouse strains (7 highly inbred) are shown in (B) and (C), respectively.

The signature of homogenization should be even stronger within cells (Stage I). The evolutionary rate of neutral rRNA variants within cells is measured in the absence of chromosome segregation and assortment. In one experiment (unpublished), the homogenization effects in rDNAs are measured in cultured cell lines derived from a single ancestral cell over 6 months of evolution. Another experiment analyzes the evolution of rRNA genes within solid tumors. We could estimate the rate at which rRNA variants spread among copies within the cells. The measurements suggest that the fixation time of new rRNA mutations within cells would take only 1 - 3 kyrs (thousand years). Since a new mutation in single-copy genes would take > 600 kyrs to be fixed in human populations, the speed of genetic drift in Stage I evolution is orders faster than in Stage II.

#### 2. rDNA divergence between species

The *C*^*\**^ values derived from the polymorphism data would have missed rapidly fixed variants, thus resulting in the under-estimation of genetic drift. We now consider the evolution of rRNA genes between species by analyzing the fixation rate of mutations. Eq. (3) shows that the mutation rate, *μ* determines the long-term evolutionary rate, *λ*. As modeled in Fig. 2, Eq. (3) is valid only when the divergence time (*T*_*d*_) between species is at least an order of magnitude longer than the fixation time (*T*_*f*_). We thus compare human and macaque monkey for this purpose; the latter is used in this study only for that purpose. The *T*_*d*_ value is 35 Myrs, whereas the *T*_*f*_ in humans is less than one Myr. As shown in Table S3, *λ* falls in the range of 50 - 60 (differences per kb) for single-copy genes and 40 - 70 for the non-functional parts of rRNA genes. The data thus suggest that rRNA and single-copy genes are comparable in mutation rate. We now analyze closely related species for the strength of homogenization in rRNA genes.

##### 1) Between mouse species – Rapid genetic drift via strong intra-individual homogenization

The four species used in this study are shown in Fig. 3A. In comparing *M. m. domesticus* with *M. spretus* and *M. m. castaneus*, there are two depths in the phylogeny with two *T*_*d*_’s (52, 53). The fourth species, *M. m. musculus* yields very similar results as *M. m. domesticus* in these two comparisons (see Supplemental Tables S4 - S5).

In Table 1, the rRNA genes are compared with the single copy genes. The rRNA genes comprise 3 different structures: the mature rRNA (18S, 5.8S and 28S), the non-functional parts of the rRNA gene (ITS and ETS) and the intergenic sequences (IGS). The mature portion is selectively constrained and is not used. The IGS region exhibits structural variability-mainly due to expansions and contractions of tandem R and long GT-rich repeats that reduce the alignment reliability (20, 39). We shall thus focus on the evolutionary rate of ETS/ITS regions in the comparison with the single-copy genes (the last two rows of Table 1).

**Table 1.** Divergence in rRNA genes at two levels of taxonomic rank.

|  |  | *M. m. domesticus* vs<br>*M. m. castaneus*<br>at the sub-species rank |  |  | *M. m. domesticus* vs<br>*Mus spretus*<br>at the species rank |  |  |
| --- | --- | --- | --- | --- | --- | --- | --- |
|  | Length<br>(bp) | Mapping<br>Length, L | Number of<br>Divergent<br>sites, D | D/L per kb | Mapping<br>Length, L | Number of<br>Divergent<br>sites, D | D/L per kb |
| 18S+5.8S + 28S | 6757 | 6755 (5321) | 4 (4) | 0.59 (0.75) | 12 (10) | 6757 (5321) | 1.78 (1.88) |
| **Non-functional parts** |  |  |  |  |  |  |  |
| 5' ETS | 4007 | 4006 (3165) | 9 (7) | 2.25 (2.21) | 4005 (3165) | 100 (68) | 24.97 (21.48) |
| ITS1+ITS2 | 2088 | 2088 (1530) | 11 (7) | 5.27 (4.58) | 2077 (1530) | 54 (38) | 26.00 (24.84) |
| 3' ETS | 551 | 539 (371) | 1 (1) | 1.86 (2.70) | 551 (371) | 17 (13) | 30.85 (35.04) |
| IGS | 31902 | 26101 (25461) | 45 (36) | 1.72 (1.41) | 25516<br>(24796) | 270 (241) | 10.58 (9.72) |
| **Single-copy genes** | - | - | - | 9.5 | - | - | 21 |
| rRNA genes<br>(averages over<br>ETS+ITS portions) | 6646 | 6633 (5066) | 21 (15) | 3.17 (2.96) | 6633 (5066) | 171 (119) | 25.78 (23.49) |

Numbers in parentheses are based on sequences with CpG sites masked.

The *M. m. domesticus* vs. *M. m. castaneus* comparisons correspond to *T*_*d1*_ of Fig. 2. At this level of divergence, rRNA gene mutations appear to be fixed more slowly (3.16/Kb) than single-copy genes (9.50/Kb). At the rank of sub-species, the divergence time (54) is close to the estimated fixation time (50, 55) and many mutations would not have been fixed. In short, the data from the two sub-species would be halfway between the data of within-species polymorphism and those of true species divergence.

Indeed, between *M. m. domesticus* and *M. spretus*, rRNA genes appear to be evolving slightly more rapidly (23.49/kb) than single-copy genes (21.0/kb) after removing all CpG sites. The ratio of 21.0/23.49 would lead to the estimation of *C*^*\**^ < 1. These results are not consistent with the projection of Fig. 2 that assumes *C*^*\**^ *≥* 1. This paradox of having fewer than one “effectively copy” can be resolved by having very high *V*^*\**^*(K)* as in *C*^*\**^ = *C/V*^*\**^*(K)*. As gene conversion, replication slippage and unequal crossover can be frequent but also highly variable in occurrences, high *V*^*\**^*(K)* is expected.

Lastly, IGS may experience fewer events of homogenization such as gene conversion and unequal crossover, leading to lower *C*^*\**^ reduction. We note that the heterozygosity in IGS region, at about 2-fold higher than that of ETS and ITS regions (8‰ for IGS, 5‰ for ETS and 3‰ for ITS in mice), supports this interpretation (see Supplemental Tables S6).

##### 2) Between Human and Chimpanzee – Beyond genetic drift in rDNA divergence

As shown in Table 2, the evolutionary rate of rRNA genes between human and chimpanzee is substantially higher than that of single-copy genes. While ITS1 and ITS2 have evolved at nearly the same rate as single copy genes, both at ~ 11 changes per kb, the bulk of the rRNA gene region has evolved at least 80% faster (~20 per kb). Hence, *C*^*\**^ < 1. However, rapid genetic drift cannot fully account of the rapid evolution. After all, even the extremely rapid fixation by drift can only increase the substitution rate by 15% [*T*_*f*_ /(*T*_*d*_ –*T*_*f*_) ~ 0.8/(6-0.8); see Fig. 2], compared to single-copy genes. An 80% increase demands other forces. Next, we will test if selection is operating on rRNA genes by examining the pattern of gene conversion.

**Table 2.**
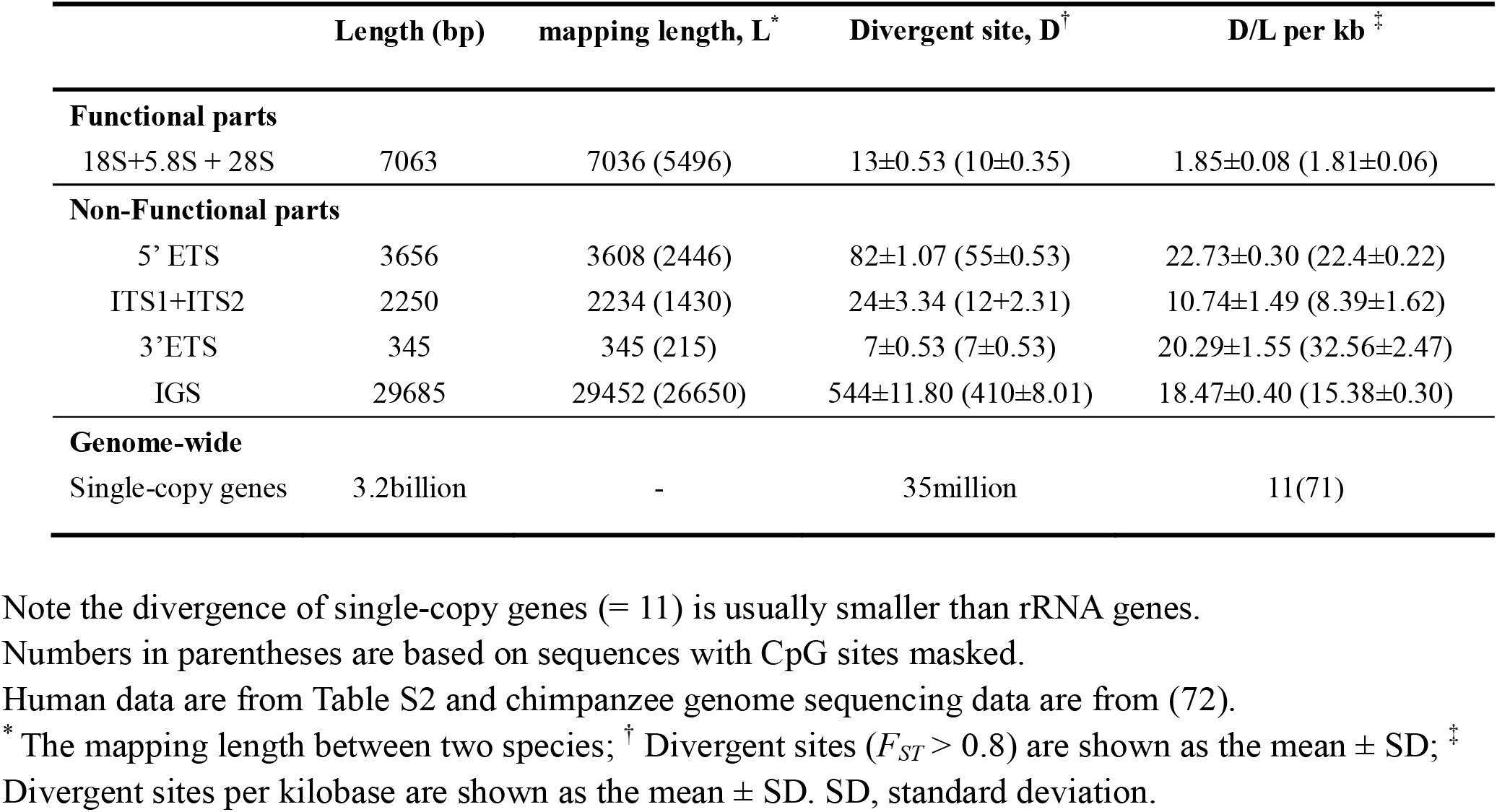
Divergence in rRNA genes between Human and Chimpanzee.

|  | Length (bp) | mapping length, L<sup>*</sup> | Divergent site, D<sup>†</sup> | D/L per kb<sup>‡</sup> |
| --- | --- | --- | --- | --- |
| **Functional parts** |  |  |  |  |
| 18S+5.8S + 28S | 7063 | 7036 (5496) | 13±0.53 (10±0.35) | 1.85±0.08 (1.81±0.06) |
| **Non-Functional parts** |  |  |  |  |
| 5' ETS | 3656 | 3608 (2446) | 82±1.07 (55±0.53) | 22.73±0.30 (22.4±0.22) |
| ITS1+ITS2 | 2250 | 2234 (1430) | 24±3.34 (12±2.31) | 10.74±1.49 (8.39±1.62) |
| 3'ETS | 345 | 345 (215) | 7±0.53 (7±0.53) | 20.29±1.55 (32.56±2.47) |
| IGS | 29685 | 29452 (26650) | 544±11.80 (410±8.01) | 18.47±0.40 (15.38±0.30) |
| **Genome-wide** |  |  |  |  |
| Single-copy genes | 3.2billion | - | 35million | 11(71) |

Note the divergence of single-copy genes (= 11) is usually smaller than rRNA genes.
Numbers in parentheses are based on sequences with CpG sites masked.
Human data are from Table S2 and chimpanzee genome sequencing data are from (72).
<sup>\*</sup> The mapping length between two species; <sup>†</sup> Divergent sites ( $F_{ST} > 0.8$ ) are shown as the mean ± SD; <sup>‡</sup> Divergent sites per kilobase are shown as the mean ± SD. SD, standard deviation.

#### 3. Positive selection for rDNA mutations in apes, but not in mice

This study makes an important point that positive selection is plausible only when genetic drift has been rigorously defined, as is done above between humans and chimpanzees. Here, a possible mechanism is selection that biases gene conversion. We hence examine the patterns of AT-to-GC vs. GC-to-AT changes. While it has been reported that gene conversion would favor AT-to-GC over GC-to-AT conversion (56, 57) at the site level, we are interested at the gene level by summing up all conversions across sites. We designate the proportion of AT-to-GC conversion as *f* and the reciprocal, GC-to-AT, as *g*. Both *f* and *g* represent the proportion of fixed mutations between species (see Methods). So defined, *f* and *g* are influenced by the molecular mechanisms as well as natural selection. The latter may favor a higher or lower GC ratio at the genic level between species. As the selective pressure is distributed over the length of the gene, each site may experience rather weak pressure.

Let *p* be the proportion of AT sites and *q* be the proportion of GC sites in the gene. The flux of AT-to-GC would be *pf* and the flux in reverse, GC-to-AT, would be *qg*. At equilibrium, *pf* = *qg*. Given *f* and *g*, the ratio of *p* and *q* would eventually reach *p*/*q* = *g*/*f*. We now determine if the fluxes are in equilibrium (*pf* =*qg*). If they are not, the genic GC ratio is likely under selection and is moving to a different equilibrium.

In these genic analyses, we first analyze the human lineage (58, 59). Using chimpanzees and gorillas as the outgroups, we identified the derived variants that became nearly fixed in humans with frequency > 0.8 (Table 3). The chi-square test shows that the GC variants had a significantly higher fixation probability compared to AT. In addition, this pattern is also found in chimpanzees (p < 0.001). In *M. m. domesticus*, the chi-square test reveals no difference in the fixation probability between GC and AT (p = 0.957). Further details of direction of fixation among each species can be found in Supplementary Figure 2. Overall, a higher fixation probability of the GC variants is found in human and chimpanzee, whereas this bias is not observed in mice.

**Table 3.** The A/T to G/C and G/C to A/T changes in apes and mouse.

|  | Direction of changes | Not fixed mutation counts | Fixed mutation counts | Chi-square test |
| --- | --- | --- | --- | --- |
| **Human** | A/T to G/C | 163 | 223 | P = 1.559e-15 |
|  | G/C to A/T | 159 | 48 |  |
| **Chimpanzee** | A/T to G/C | 289 | 210 | P = 0.000469 |
|  | G/C to A/T | 201 | 83 |  |
| ***M. m. domesticus*** | A/T to G/C | 99 | 10 | P = 0.9568 |
|  | G/C to A/T | 156 | 14 |  |

Based on Table 3, we could calculate the value of *p, q, f* and *g*. Shown in the last row of Table 4, the (*pf*)/(*qg*) ratio is much larger than 1 in both the human and chimpanzee lineages. Notably, the ratio in mouse is not significantly different from 1. Combining Tables 2 and 4, we conclude that the slight acceleration of fixation in mice can be accounted for by genetic drift, due to homogenization among rRNA gene copies. In contrast, the different fluxes corroborate the interpretations of Table 2 that selection is operating in both humans and chimpanzees.

**Table 4.** The parameter values of *p, q, f* and *g* in the evolution between A/T and G/C.

| Species | Human |  |  | Chimpanzee |  |  | *M. m. domesticus* |  |  |
| --- | --- | --- | --- | --- | --- | --- | --- | --- | --- |
| Sequence segments | Non-functiona<br>l | Total<br>rDNA | Whole<br>genome | Non-functional | Total<br>rDNA | Whole<br>genome | Non-functional | Total<br>rDNA | Whole<br>genome |
| $p$ | 0.43 | 0.42 | | 0.43 | 0.42 | | 0.54 | 0.51 | |
| $q$ | 0.57 | 0.58 | 0.41 | 0.57 | 0.58 | 0.42 | 0.46 | 0.49 | 0.42 |
| $f$ | 0.0142 | 0.0125 | | 0.0136 | 0.012 | | 0.0005 | 0.0004 | |
| $g$ | 0.0023 | 0.0019 | | 0.0041 | 0.0034 | | 0.0007 | 0.0006 | |
| $(pf)/(qg)$ | 4.66 | 4.76 | | 2.5 | 2.56 | | 0.84 | 0.69 | |

## Discussion

The Haldane (or GH) model is an “individual-output” model of genetic drift (1, 45). Hence, it does not demand the population to follow the rules of WF populations. Indeed, it would be difficult to define WF populations in any multi-copy gene systems. The two tiers of populations experience very different random forces (homogenization within individuals and conventionl drift between individuals) that together account for the genetic drift in such systems.

There have been many rigorous analyses that confront the homogenizing mechanisms directly. These studies (24–28) modeled gene conversion and unequal cross-over head on. Unfortunately, on top of the complexities of such models, the key parameter values are rarely obtainable. In the branching process, all these complexities are wrapped into *V*^*\**^(*K*) for formulating the evolutionary rate. In such a formulation, the *collective* strength of these various forces may indeed be measurable. By compressing all stochastic elements into *V(K)*, or *V*^*\**^*(K)*, we will be able to filter out random factors and focus on forces of greater biological interest such as selection. Interestingly, while rDNA evolution in mice can be fully accounted for by genetic drift, its evolution in the great apes must have been driven by selection along with genetic drift. The contrast shows that the GH model can account for genetic drift more comprehensively than the WF models.

The branching process is a model for general processes. Hence, it can be used to track genetic drift in most systems with two stages of evolution, even though TEs, viruses and rRNA genes are very different biological entities. In multi-copy genes, like rDNA, the drift is strong enough to reduce the copy number in the entire population from ~ 150*N* to < 1*N*. This acceleration is seen in mice but would have been interpreted to be due to positive selection in the conventional analyses. Nevertheless, positive selection is indeed evident in human and chimpanzee, as the evolutionary rate exceeds the limit of strong drift.

In conclusion, the GH model is far more general than the WF model. Its *E*(*K*) parameter would define *N*’s while *V*(*K*) would track random changes in gene frequency. The chaotic analyses of COVID-19 have shown why an accurate account of genetic drift is indispensable (14, 23, 29, 60). In this study, we use the rRNA gene system for such a demonstration.

## Materials and Methods

### Data Collection

We collected high-coverage whole-genome sequencing data for our study. The genome sequences of human (n = 8), chimpanzee (n = 1) and gorilla (n = 1) were sourced from National Center for Biotechnology Information (NCBI) (Table S7). Human individuals were drawn from diverse geographical origins encompassing three continents (4 Asians, 2 Europeans and 2 Africans) and Asia CRC was the normal tissue of Case 1 patient from (12).

Genomic sequences of mice (n = 13) were sourced from the Wellcome Sanger Institute’s Mouse Genome Project (MGP) (61). Although some artificial selection has been performed on laboratory mouse strains (62), the WSB/EiJ, ZALENDE/Ei and LEWES/EiJ strains were derived from wild populations. Incorporating these wild-derived laboratory strains, along with other inbred strains, a cohort of 10 mice was utilized to approximate the population representation of *M. m. domesticus*. Furthermore, the low *F*_*IS*_ of 0.14 for rDNA in *M. m. domesticus* found in this study suggests that each mouse covers 86% of the population’s genetic diversity, thereby mitigating concerns about potential sampling biases.

Accessions and the detailed information of samples used in this study are listed in the Table S7 and Table S8.

### Variant allele frequency

Following adapter trimming and the removal of low-quality sequences, these whole-genome sequencing data of apes and mice were mapped against respective reference sequences: the human rDNA reference sequence (Human ribosomal DNA complete repeating unit, GenBank: U13369.1) and the mouse rDNA reference sequence (*Mus musculus* ribosomal DNA, complete repeating unit, GenBank: BK000964.3). Alignment was performed using Burrows-Wheeler-Alignment Tool v0.7.17 (63) with default parameters. All mapping and analysis are performed among individual copies of rRNA genes.

Each individual was considered as a psedo-population of rRNA genes and the diversity of rRNA genes was calculated using this psedo-population of rRNA genes. To determine variant frequency within individual, variants were called from multiple alignment files using bcftools (64) to ensure the inclusion of all polymorphic sites that appeared in at least one sample and to maintain consistent processing steps. Per-sample variant calling results were generated in a VCF file using the following settings: ‘bcftools mpileup --redo-BAQ --max-depth 50000 --per-sample-mF --annotate’ and ‘bcftools call -mv’.

Our analysis specifically focused on single nucleotide variants (SNVs) with only two alleles, while other mutation types were discarded. Variant information of interest per sample was extracted from the VCF files using the command ‘bcftools view -V indels’ and

‘bcftools query –f ‘%CHROM\t%POS\t%REF\t%ALT\t[\t%GT]\t[\t%DP]\t[\t%AD]\n’\’.

The AD (total high-quality bases of allelic depths) was used to obtain the number of reference-supporting reads (n.ref) and alternative-supporting reads (n.alt) in each sample. Sites with a depth < 10 in any sample were filtered out, resulting in an average of the minimum depths above 3000 for each remaining site across all samples. This filtering step enhances the robustness of variant frequency estimation, where the variant frequency was approximated by the ratio n.alt/(n.ref + n.alt). This process allowed us to identify all polymorphic sites present in the samples and the variant frequency within each individual. Then the population-level variant frequency was computed by averaging variant frequencies across all individuals.

### Identification of Divergence Sites

Polymorphism represents the transient phase preceding divergence. With variant frequency within individual, within species, and between species, we obtained the *F*_*IS*_ and *F*_*ST*_ values for each site. *F*_*ST*_ analysis between human and chimpanzee was conducted for each human individual, summarized in Table 2. We identified a range from 672 to 705 sites with *F*_*ST*_ values above 0.8 across individuals, depicting robust divergence sites. Considering the high mutation rate in CpG sites (65, 66) and predominantly GC content in rDNA (Table 4), we further estimated the evolutionary rate at non-CpG sites during the interspecies divergence. To achieve this, mutations in CpG sites were manually removed by excluding all sites containing CpG in one species and TpG or CpA in the other; the reverse was similarly discarded. Additionally, the count of non-CpG sites within the mapping length, where site depth exceeded 10, was performed by Samtools (64) with the settings ‘samtools mpileup -Q 15 -q 20’. As a result, the evolutionary rate of rDNA in non-CpG sites was ascertained.

For assessing diversity and divergence across gene segments, we used ‘samtools faidx’ to partition variants into a total of 8 regions within rRNA genes, including 5’ETS, ITS1, ITS2, 3’ETS, IGS, 18S, 5.8S, and 28S, aligning them with corresponding reference sequences for further analysis. The functional parts (18S, 5.8S, and 28S) were subject to strong negative selection, exhibiting minimal substitutions during species divergence as expected. This observation is primarily used for comparison with the non-functional parts.

### Genome-wide Divergence Estimation

To assess the genome-wide divergence between 4 mouse strain species, we downloaded their toplevel reference genomes from Ensembl genome browser (GenBank Assembly ID: GCA_001624865.1, GCA_001624775.1, GCA_001624445.1, and GCA_001624835.1). Then we used Mash tools (67) to estimate divergence across the entire genome (mainly single-copy genes) with ‘mash sketch’ and ‘mash dist’. Additionally, the effective copy number of rRNA genes, denoted as *C*^*\**^, can be estimated by calculating the ratio of population diversity observed in rDNA to that observed in single-copy genes.

### Estimation of Site Conversion

We identified derived alleles in human, chimpanzees, and *M. m. domesticus* by comparing each lineage to two outgroups (chimpanzee and gorilla, humans and gorilla, *M. m. castaneus* and *M. spretus*) and defining the ancestral state as alleles present in both outgroups at frequency >0.8. This approach minimizes the influence of derived mutations’ initial frequency on fixation probability and fixation time.

Derived variants were then categorized by their frequency into two groups: (nearly) fixed (> 0.8) vs not fixed (as shown in Table 3). The frequency threshold of >0.8 was chosen to balance the need for a sufficient number of sites to calculate of (*f*/*g*), and to ensure reliability. We also applied a more stringent threshold of >0.9, which yielded similar results.

In this study, six types of mutations were tabulated, representing ancestral-to-derived as depicted in Supplementary Fig. S2. For example, A-to-G represented the both A-to-G and T-to-C types of mutations. The C-to-G (or G-to-C) and A-to-T (or T-to-A) types of mutations were excluded in the subsequent analysis.

Specifically, *f* represents the proportion of fixed mutations where an A or T nucleotide has been converted to a G or C nucleotide, normalized over total A/T sites. *g* represents the proportion of fixed mutations where an G or C nucleotide has been converted to a A or T nucleotide, normalized over total G/C sites.

The consensus rDNA sequences for the species lineage were generated by Samtools consensus (64) from the bam file after alignment. The following command was used:

‘samtools consensus -@ 20 -a -d 10 --show-ins no --show-del yes input_sorted.bam output.fa’.

Furthermore, the alternative hypotheses of GC-biased mutation process (68, 69) alone can be rejected in this study. According to the prediction of the mutational mechanism hypotheses, AT or GC variants should have equal fixation probabilities. We compared nearly fixed AT to GC versus GC to AT mutations using chi□square tests. We observed significant fixation bias favoring GC in apes, but not in mice, discounting mutation bias alone.

## Data Availability

No new data were generated in this study. The genomic data used in this study are available from National Center for Biotechnology Information (NCBI) (https://www.ncbi.nlm.nih.gov/) and the Mouse Genomes Project (https://www.sanger.ac.uk/data/mouse-genomes-project/). The specific accession numbers recorded in Supplementary Table S7 and S8.

## Acknowledgments

We are grateful for the helpful comments from many colleagues, in particular, Weiwei Zhai, Yong Zhang, GuoDong Wang of CAS and Jianrong Yang of SYSU. This work was supported by the National Natural Science Foundation of China (32150006, 32293193/32293190 to C.I.W., 32200493 to Y.R., and 82341092 to HJ Wen.), the National Key Research and Development Projects of the Ministry of Science and Technology of China (2021YFC2301300, 2021YFC0863400), and Guangdong Key Research and Development Program (No. 2022B1111030001).

