## supplementary information for "The paradox of extremely fast evolution driven by genetic drift in multi-copy gene systems"

**This PDF file includes:**

Supporting Text

Figures S1 to S2

Tables S1 to S8

SI References

**Supporting Information Text**

**Note 1. Basic theory for multi-copy genes (rDNAs) polymorphism within species**

A standard measure of genetic drift is the level of heterozygosity (H). At the mutation-selection equilibrium

$$H_{equi}=\frac{{2N}_{e}\mu}{{2N}_{e}\mu+1}$$

where *μ* is the mutation rate of the entire gene and *N_e_* is the effective population size. In this study, *N_e_* = *N* for single-copy gene and *N_e_* = *C^*^N* for rRNA genes. The empirical measure of nucleotide diversity *H* is given by

$$H=\frac{\sum_{i=1}^{L} 2p_{i}(1-p_{i})}{L} Eq. (2)$$

where *L* is the gene length (for each copy of rRNA gene, *L* ~ 43kb) and *p_i_* is the variant frequency at the *i*-th site. As rRNA genes exist as paralogs rather than orthologs, the diversity refers to the sequence variation across all rDNA copies within the genome. This diversity can be assessed at both individual and population levels.

We calculate *H* of rRNA genes at three levels – within-individual, within-species and then, within total samples (H_I_, H_S_ and H_T_, respectively). *H_S_* and *H_T_* are standard population genetic measures (1, 2). In calculating *H_S_*, all sequences in the species are used, regardless of the source individuals. A similar procedure is applied to *H_T_*_._ The *H_I_* statistic is adopted for multi-copy gene systems for measuring within-individual polymorphism. Note that copies within each individual are treated as a pseudo-population (see Fig. 1). With multiple individuals, *H_I_* is averaged over them.

Given the three levels of heterozygosity, there are two levels of differentiation. First, *F_IS_* is the differentiation among individuals within the species, defined by

*F_IS_ = [H_S_ - H_I_]/H_S_*

*F_IS_* is hence the proportion of genetic diversity in the species that is found only between individuals. We will later show *F_IS_ ~* 0.05 in human rDNA (Table S2), meaning 95% of rDNA diversity is found within individuals.

Second, *F_ST_* is the differentiation between species within the total species complex, defined as

*F_ST_ = [H_T_ – H_S_]/H_T_*

*F_ST_* is the proportion of genetic diversity in the total data that is found only between species. *F_ST_* closed to 1 would indicate a large genetic distance between species.

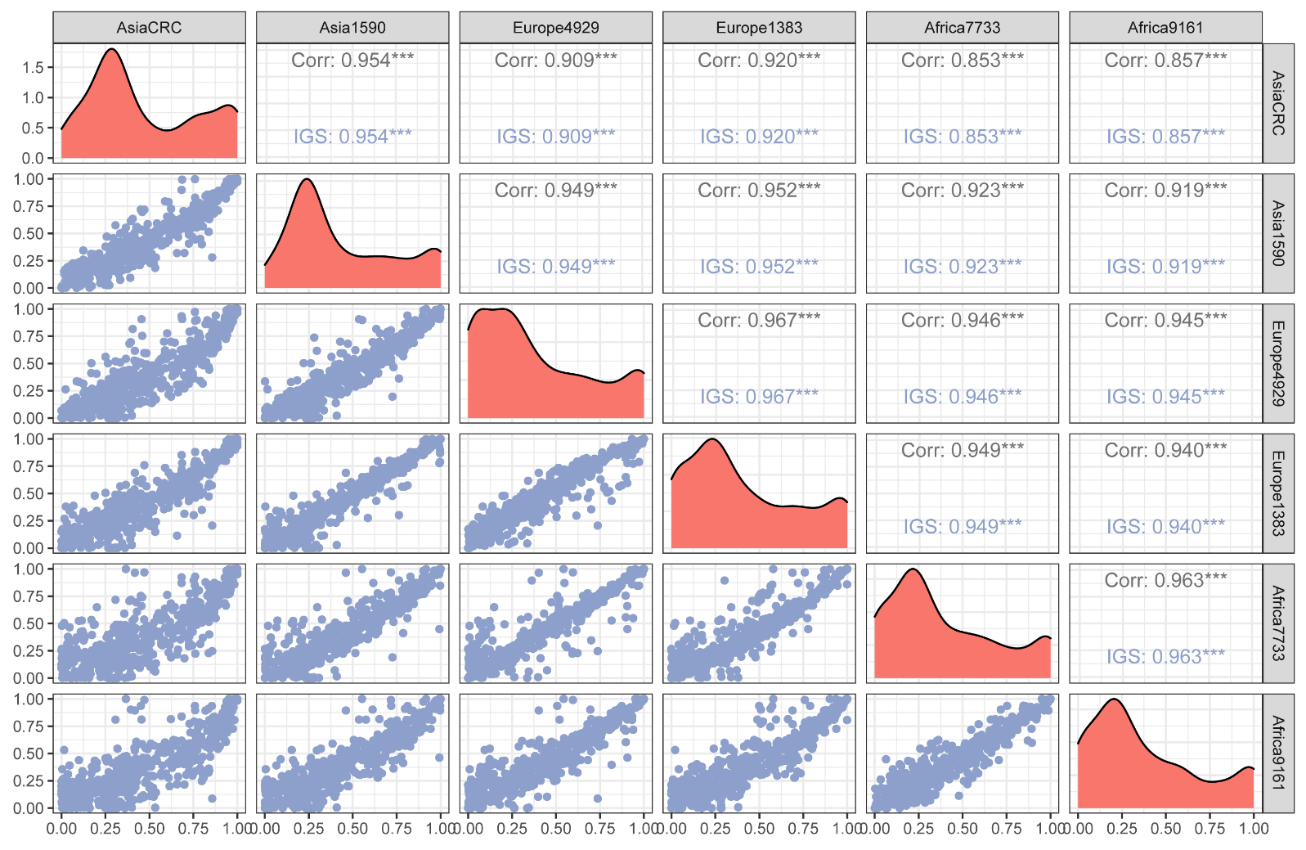

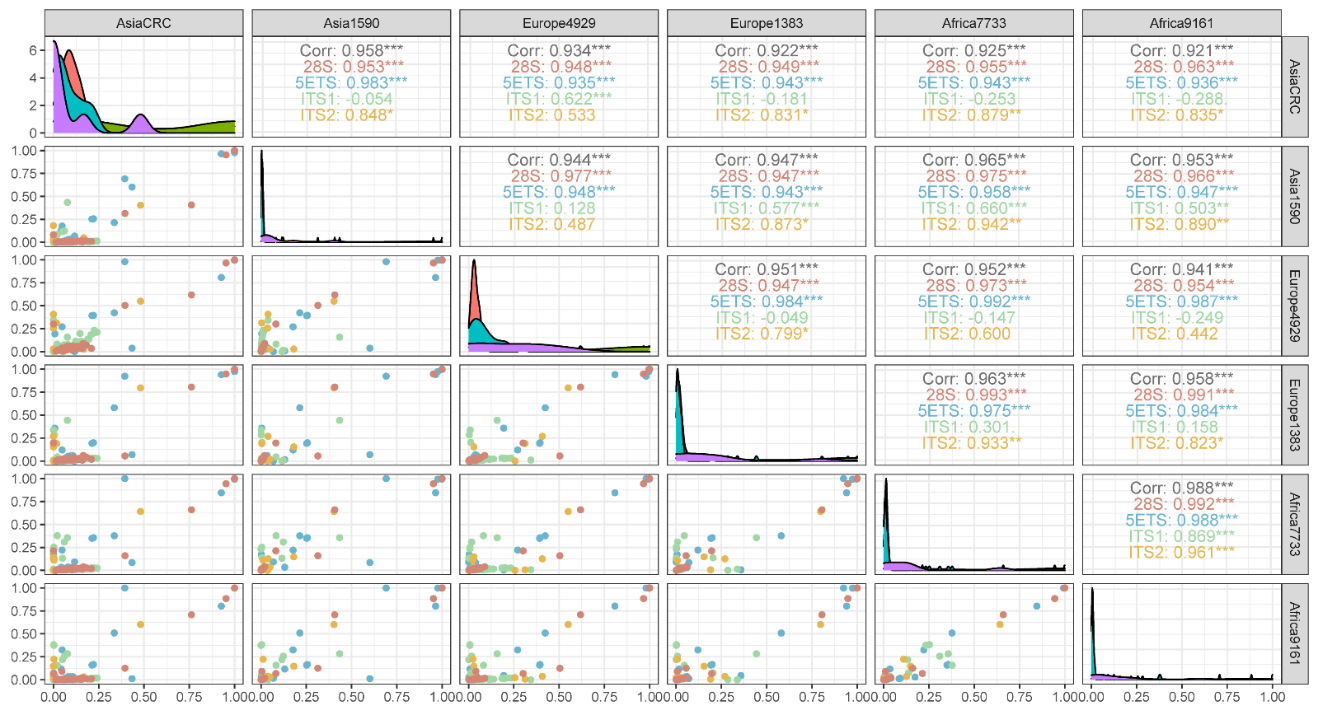

**Figure S1. Correlation of variant frequencies between human individuals.** The pairwise correlation of variant site frequency in the transcribed region(top) and IGS region (bottom) of rDNAs among 6 individuals (2 Asians, 2 Europeans, and 2 Africans). The high inter-individual correlations suggest that the rDNA diversity is mainly preserved within each individual. Each color represents a region of rDNA. The diagonal plots present the variant frequency distribution. The upper right section summarizes the Pearson correlation coefficients derived from the mirror symmetric plots in the bottom left panels. The analysis excluded the 18S, 3'ETS, and 5.8S regions due to the limited polymorphic sites.

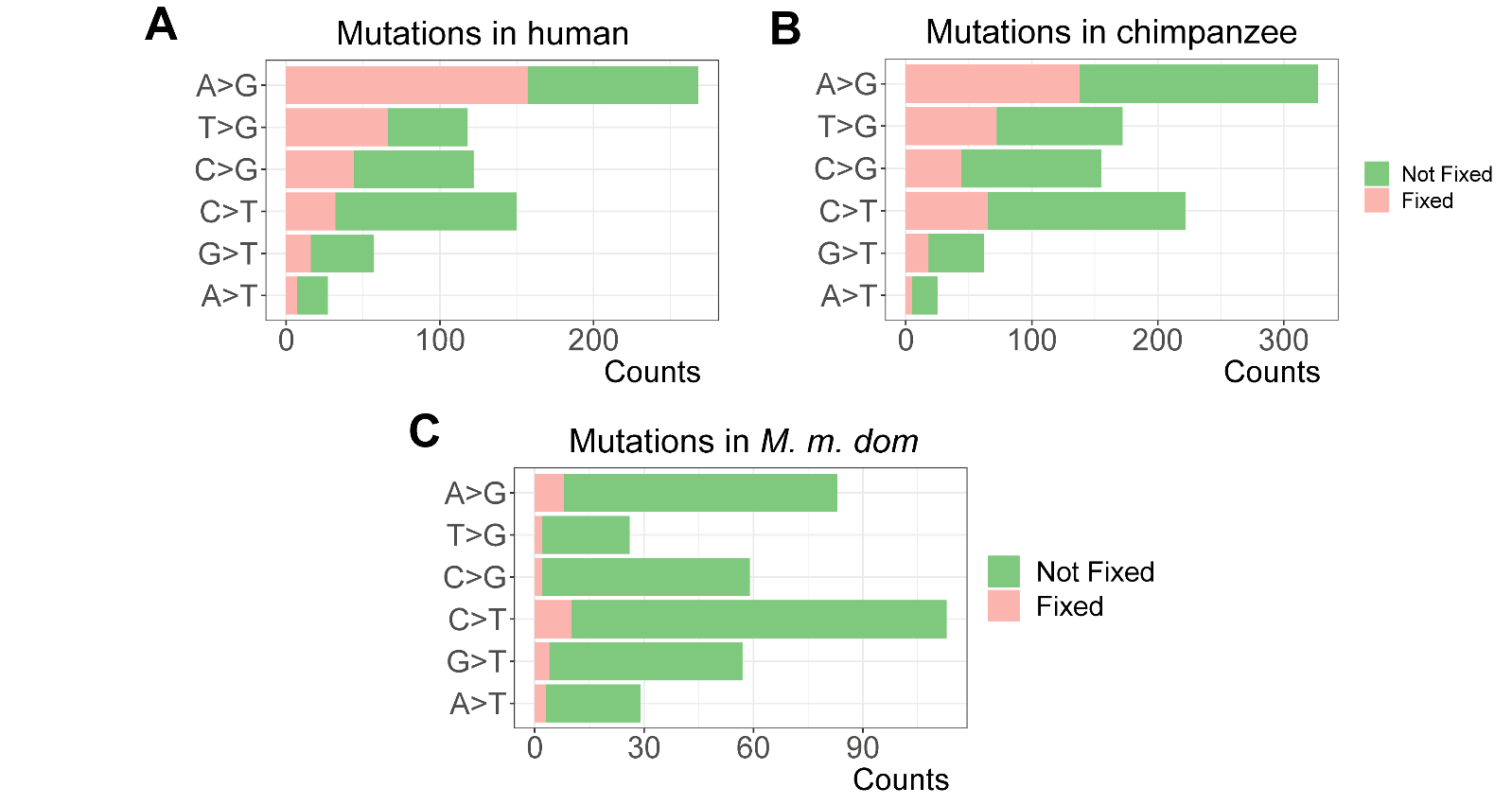

Fig. S2. The different fixation proportion of mutation in apes and mouse. Mutations are divided into 6 types and shown in the figure for the human (A), chimpanzee (B), and *M. m. domesticus* (C) lineages, respectively. The histogram is filled with red color to represent the nearly fixed proportion, while the green color represents the remaining proportion (not fixed). Biased GC fixation probability is found in human and chimpanzee, but not in *M. m. domesticus*.

**Table S1.** rRNA gene nucleotide diversity in the 10 *M. m. domesticus* strains of a global collection.

| **Mouse strains** | ***H_I_* (‰) - diversity within individuals** | |
| --- | --- | --- |
|  | **Functional parts**  **(18S 5.8S and 28S)** | **Non-functional parts**  **(ETS, ITS and IGS)** |
| WSB/EiJ | 0.36 | 6.45 |
| ZALENDE/Ei | 0.44 | 6.29 |
| LEWES/EiJ | 0.28 | 5.56 |
| BALB/cJ | 0.19 | 6.51 |
| C57BL/6NJ | 0.20 | 6.30 |
| ST/bJ | 0.24 | 6.73 |
| NZW/LacJ | 0.22 | 4.96 |
| FVB/NJ | 0.35 | 6.47 |
| DBA/1J | 0.22 | 6.44 |
| C3H/HeJ | 0.20 | 6.50 |
| Averaged diversity ***H_I_*** across individuals | 0.27 | **6.22** |
| ***H_S_* for rDNA in *M. m. domesticus*** | 0.34 | 7.25 |
| ***H_S_* for single-copy genes in *M. m. domesticus*** | --- | 1.40^*^ |

WSB/EiJ, ZALENDE/Ei and LEWES/EiJ are outbred strains, and the rest are inbred strains.

Sources: Wellcome Sanger Institute’s Mouse Genome Project (MGP).

^*^Data from (3).

**Table S2.** rRNA gene nucleotide diversity in the 8 humans of a global collection.

| **Human Individuals** | ***H_I_* (‰) - diversity within individuals** | |
| --- | --- | --- |
|  | **Functional parts**  **(18S 5.8S and 28S)** | **Non-functional parts**  **(ETS, ITS and IGS)** |
| Asia CRC | 0.53 | 7.69 |
| Asia 1590 | 0.28 | 7.33 |
| Europe 4929 | 0.42 | 6.53 |
| Europe 1383 | 0.32 | 6.74 |
| Africa 7733 | 0.36 | 6.65 |
| Africa 9161 | 0.34 | 6.55 |
| Asia F9551 | 0.87 | 6.37 |
| Asia M9552 | 0.88 | 6.58 |
| Averaged diversity ***H_I_*** across individuals | 0.50 | 6.81 |
| ***H_S_* for rDNA in human population** | 0.68 | 7.24 |
| ***H_S_* for single-copy genes in human population** | --- | 0.88^*^ |

8 humans are from three continents (4 Asians, 2 Europeans and 2 Africans).

Source: National Center for Biotechnology Information (NCBI).

^*^ Data from (4).

Table S3. Divergence in rRNA genes between Human and Rhesus Macaque.

|  | **Length** | **mapping length L^*^**  (w/o CpG sites) | **Divergent site D**  (w/o CpG sites) | **D/(L/1000)** ^†^  (w/o CpG sites) |
| --- | --- | --- | --- | --- |
| **Functional parts** | | | | |
| 18S+5.8S + 28S | 7063 | 6824(5364) | 28 (16) | 4.10 (2.98) |
| **Non-Functional parts** | | | | |
| 5’ ETS | 3656 | 2586(1898) | 180 (104) | 69.61 (54.79) |
| ITS1+ITS2 | 2250 | 1762(1182) | 68 (54) | 38.59 (45.69) |
| 3’ETS | 345 | 295(199) | 15 (11) | 50.85 (55.28) |
| IGS | 29685 | 24254(22722) | 946 (703) | 39.00 (30.94) |
| **Genome-wide** | | | | |
| Single-copy genes |  |  |  | 50-60 (5-7) |

^*^ The mapping length between two species; ^†^ Divergent sites per kilobase are shown.

Table S4. Divergence sites in rRNA genes between *M. m. musculus* and *M. m. castaneus*.

|  | **Length** | **mapping length L**  **(w/o CpG sites)** | **Divergent site D**  **(w/o CpG sites)** | **D/(L/1000)**  **(w/o CpG sites)** |
| --- | --- | --- | --- | --- |
| **Functional parts** | | | | |
| 18S+5.8S + 28S | 6757 | 6757(5321) | 2(2) | 0.3(0.4) |
| **Non-Functional parts** | | | | |
| 5’ ETS | 4007 | 4005(3165) | 18(11) | 4.5(3.5) |
| ITS1+ITS2 | 2088 | 2077(1530) | 2(2) | 1.0(1.3) |
| 3’ETS | 551 | 551(371) | 1(1) | 1.8(2.7) |
| IGS | 31902 | 26233(25483) | 36(29) | 1.4(1.1) |
| **Genome-wide** | | | | |
| Single-copy genes | - | - | - | 9.9 |

Table S5. Divergence sites in rRNA genes between *M. m. musculus* and *Mus spretus*.

|  | **Length** | **mapping length L**  **(w/o CpG sites)** | **Divergent site D**  **(w/o CpG sites)** | **D/(L/1000)**  **(w/o CpG sites)** |
| --- | --- | --- | --- | --- |
| **Functional parts** | | | | |
| 18S+5.8S + 28S | 6757 | 6757(5321) | 13(10) | 1.9(1.9) |
| **Non-Functional parts** | | | | |
| 5’ ETS | 4007 | 4005(3165) | 106(70) | 26.5(22.1) |
| ITS1+ITS2 | 2088 | 2077(1530) | 57(41) | 27.4(26.8) |
| 3’ETS | 551 | 551(371) | 19(15) | 34.5(40.4) |
| IGS | 31902 | 25516(24796) | 279(249) | 10.9(10.0) |
| **Genome-wide** | | | | |
| Single-copy genes | - | - | - | 21 |

The divergence between *M. m. musculus* and the two outgroups species of *M. m. castaneus* and *Mus spretus* are shown. These findings are similar to the results in *M. m. domesticus*.

**Table S6.** Average nucleotide diversity (‰) in population (*H_S_*)

|  | functional region | IGS | ETS | ITS | non-functional region |
| --- | --- | --- | --- | --- | --- |
| *Mus. m. dom* (n=10) | 0.34 | 7.95 | 5.13 | 3.27 | 7.25 |
| Human  (n=8) | 0.68 | 8.16 | 2.8 | 3.12 | 7.24 |

Table S7. Sample information of apes.

| **Sample** | **Sample information** | **Accession** | **Submitted by** |
| --- | --- | --- | --- |
| Asia CRC | 70-year-old  female Asian | Normal pair of Case 1 | Chen B, *et al*. (2022)(8) |
| Asia 1590 | 60-year-old  female Asian | SRR13921590 | National Cheng Kung University Hospital |
| Asia F9551 | 44-year-old  male Asian | SAMN03009551 | VNU University of Engineering and Technology |
| Asia M9552 | 40-year-old  female Asian | SAMN03009552 | VNU University of Engineering and Technology |
| Europe 4929 | 51-year-old  male European | SRR11184929 | Garvan Institute of Medical Research |
| Europe 1383 | male  European | HGDP01383 | The Wellcome Trust Sanger Institute (SC) |
| Africa 7733 | male  African | HGDP00985 | The Wellcome Trust Sanger Institute (SC) |
| Africa 9161 | male  African | HGDP00461 | Harvard Medical School, Department of Genetics |
| Chimpanzee | male  Pan troglodytes | SAMD00026331 | National Institute of Genetics (Japan) (9) |
| Gorilla | female  Western lowland gorilla | SAMN01920478 | IBE (CSIC-Universitat Pompeu Fabra) |

Table S8. Sample information of Mice from Mouse Genomes Project.

| **Strain** | **Accession** | **Taxa** | **Classical inbred** |
| --- | --- | --- | --- |
| WSB/EiJ | ERR9882525 | *Mus musculus domesticus* | FALSE |
| ZALENDE/Ei | ERR9882529 | *Mus musculus domesticus* | FALSE |
| LEWES/EiJ | ERR9880679 | *Mus musculus domesticus* | FALSE |
| BALB/cJ | ERR9880211 | *Mus musculus domesticus* | TRUE |
| C57BL/6NJ | ERR9880493 | *Mus musculus domesticus* | TRUE |
| ST/bJ | ERR9881367 | *Mus musculus domesticus* | TRUE |
| NZW/LacJ | ERR9880710 | *Mus musculus domesticus* | TRUE |
| FVB/NJ | ERR9880654 | *Mus musculus domesticus* | TRUE |
| DBA/1J | ERR9880632 | *Mus musculus domesticus* | TRUE |
| C3H/HeJ | ERR9880473 | *Mus musculus domesticus* | TRUE |
| CAST/EiJ | ERR9880601 | *Mus musculus castaneus* | FALSE |
| PWK/PhJ | ERR9880720 | *Mus musculus musculus* | TRUE |
| SPRET/EiJ | ERR9880927 | *Mus spretus* | FALSE |
